# The impact of language proficiency on task-dependent neural activity and functional connectivity: Insights from deafness

**DOI:** 10.1101/2024.08.30.609857

**Authors:** Valeria Vinogradova, Barbara Manini, Bencie Woll, Martin Eimer, Velia Cardin

## Abstract

Our study aimed to investigate the impact of differences in language proficiency on the neural correlates of cognitive processing in deaf adults. In congenitally and early deaf individuals, individual differences in language proficiency reflect the degree of language access during critical developmental periods and significantly influence cognitive function. By studying the neural substrates of cognition in a population with diverse language backgrounds and skills, we can explore the influence of early language experience on the formation of cognitive networks in the brain. We used functional MRI in a group of deaf adults with varying language experience backgrounds and a control group of hearing participants. In this study, we investigated the hypothesis that differences in language skills modulate neural response and functional connectivity during the execution of demanding cognitive tasks tapping into working memory and planning. Our study revealed that differences in language proficiency, independently of language modality (signed or spoken), are positively correlated with neural activity and functional connectivity within regions of the task-positive network during working memory in deaf adults. Furthermore, compared to hearing participants, the deaf group showed distinctive patterns of neural activity and connectivity during working memory task performance in task-dependent regions of the brain. Taken together, our findings emphasise the profound impact of early environmental experiences on brain responses during cognitive processing. Specifically, they highlight the role of language proficiency in shaping and supporting high-order cognition in the brain.

**Significance statement:** Higher-order cognitive abilities, such as working memory and planning, are essential for human activity. They enable the coordination of mental processes and goal-directed behaviour. When successfully executing cognitive tasks, the human brain exhibits systematic patterns of neural activity and connectivity in the frontal and parietal regions. In this study, we investigated whether the control of nonverbal cognition is influenced by differences in language proficiency at the level of the functional organisation of the brain.

Our study showed that differences in language proficiency in deaf individuals are positively associated with neural activity and functional connectivity within the task-positive brain regions during working memory. These findings offer compelling evidence for the critical role of access to a natural language in scaffolding the neural mechanisms that support higher-order cognition. Our results provide a deeper understanding of the relationship between language proficiency and cognitive processing, highlighting the importance of early language access in modulating and shaping brain function.

Revealing the contribution of a developed language system to the formation of cognitive networks has implications not only for understanding the processes underlying cognitive development and functioning but also for informing policy and practice in relation to education and language access.

## Introduction

The relationship between language and cognition is one of the central foci of inquiry in psychology and neuroscience. Individuals with atypical language and communication experiences often show differences in cognitive skills, namely executive function (EF), when compared to typically developing peers (developmental language disorder: Blom & Boerma, 2020; Henry et al., 2012; Im-Bolter et al., 2006; Marini et al., 2020; Pauls & Archibald, 2016; autism: Bishop & Norbury, 2005; Demetriou et al., 2018; Weismer et al., 2018). Associations between language and cognitive function have also been noted in typically developing children (Kaushanskaya et al., 2017; Khanna & Boland, 2010; Velichkovsky et al., 2019; Weiland et al., 2014; Woodard et al., 2016). However, establishing direct causation between the two domains remains challenging due to inconsistent and bidirectional evidence of associations in different populations (Akbar et al., 2013; Friedman & Sterling, 2019; Gooch et al., 2016; Henry et al., 2012; Kuhn et al., 2014; Slot & von Suchodoletz, 2018). Investigating the effects of language proficiency on cognitive networks in the brain could uncover the precise mechanisms behind the interplay between language and cognition.

To date, research has predominantly focused on neurodivergent populations with co-occurring difficulties in language and cognitive skills. Within these groups, the differences from the typically developing peers in language and cognitive function likely stem from neural origins. Studying the neural mechanisms of cognitive function in deaf individuals allows a direct investigation of the role of language in shaping cognition in the brain. Unlike other populations that show variability in both language and cognitive skills, delays in language development in deaf children arise from limited access to language, rather than intrinsic changes in neural function. Deaf children who are born to deaf parents usually have accessible visual language input from birth from their parents.

In this way, they acquire sign language through the same developmental milestones as their hearing peers acquire spoken language (Anderson & Reilly, 2002; Mayberry & Squires, 2006; Morgan & Woll, 2002; Schick, 2003). Critically, only a minority of deaf children are born to deaf parents and are exposed to sign language from birth (less than 10%; Mitchell & Karchmer, 2004). The majority of deaf children may not have sufficient regular and meaningful engagement with an accessible language during development (Humphries et al., 2012). Despite the growing use of advanced hearing technology, such as cochlear implants, it does not ensure that a deaf child will display the same level of language attainment as their hearing peers (Boons et al., 2013; Duchesne et al., 2009; Geers et al., 2009; Lund, 2016; Tobey et al., 2013).

Delayed language acquisition in deaf children has cascading effects on other domains. EF, comprising the cognitive processes that coordinate thought in action and underline goal-directed behaviour, is particularly affected in deaf children, who have been reported to perform lower than their hearing peers on various EF tasks (Beer et al., 2014; Boerrigter et al., 2023; Botting et al., 2017; Burkholder & Pisoni, 2003; Figueras et al., 2008; Harris et al., 2013; Hintermair, 2013; Jones et al., 2020; Kronenberger et al., 2013). Critically, studies on language and EF in deafness show that reported challenges in EF can be explained by language background or skills (Botting et al., 2017; Figueras et al., 2008; Kotowicz et al., 2023; Marshall et al., 2015; Merchán et al., 2022). These findings suggest that environmental factors, such as parent-child communication and language learning (Morgan et al., 2021), but not deafness itself (Hall et al., 2018), influence language and cognitive development in deaf children.

If language experiences contribute to the formation of cognitive function during development, do differences in language skills modulate the neural substrates of higher-order cognition? While behavioural studies point to associations between deafness, language skills, and cognitive function, neuroimaging can reveal how differences in language proficiency may influence brain networks responsible for cognitive processing. Goal-directed cognition is supported by neural activity in the ‘task-positive’ (TP) regions (Fox et al., 2005) and by the interplay between this frontoparietal TP network and the default mode (or ‘task-negative’ [TN]; Buckner et al., 2008; Raichle et al., 2001; Shulman et al., 1997) network (Kelly et al., 2008; Spreng et al., 2010). These networks display antagonistic neural profiles (Cheng et al., 2020) and anticorrelated patterns of functional connectivity (Fox et al., 2005; Fransson, 2005; Kelly et al., 2008).

If the development of language skills provides the foundation for cognitive processing by modulating the neural foundations of cognitive function, neural response and functional connectivity of task-dependent brain areas will be associated with differences in language skills in individuals with varying early language experiences. In deaf individuals, language acquisition and skills are determined by access to language input early in life: the age of sign language acquisition has been linked to measures of proficiency even after decades of sign language use in this population (Cormier et al., 2012; Mayberry & Fischer, 1989; Tomaszewski et al., 2022). Deaf native signers also are more accurate in grammatical judgements of written English sentences than non-native signers, suggesting a role of timely first language exposure in language development across modalities (Mayberry, 2002). In hearing people, the association between early language experience and proficiency later in life is difficult to capture due to the homogeneity of their auditory experiences that allow natural acquisition of spoken languages. By examining how differences in language proficiency relate to neural response and functional connectivity in task-related networks in the brain, our research offers valuable insights into the neural underpinnings of cognitive function and the role of language experience in shaping cognition.

## Materials and methods

### Participants

29 congenitally or early deaf and 20 hearing individuals participated in this study. Participants in both groups were right-handed and had full or corrected-to-full vision. None of the participants had any known neurological or psychiatric conditions. Data from all participants included in this study have been previously published in a paper investigating crossmodal plasticity in the auditory cortex (Manini et al., 2022).

The information on deafness and language background was collected with a detailed questionnaire prior to the testing session. All hearing participants were native English speakers. Deaf participants had either English or British Sign Language (BSL)^1^ as their first language (Table 1). The data collection protocol included audiometric screening for deaf participants. Alternatively, deaf participants could provide copies of audiometric reports. One deaf participant was excluded from data analysis because the audiometric testing indicated a mild degree of deafness (pure-tone average in the best ear: < 25 db). Data from two participants were excluded from both of the experiments reported in this article because their performance was at chance in the conditions of interest. Data from three more participants were excluded from the analysis of the planning experiment due to excessive motion, not performing the task, or misunderstanding the rules. In total, data from 24 deaf and 20 hearing participants were included (working memory: n_deaf_ = 23, n_hearing_ = 19; planning: n_deaf_ = 20; n_hearing_ = 19). The final groups of participants were matched on age (range_deaf_ = 19-66; M ± SD_deaf_ = 40.96 ± 14.21; range_hearing_ = 18-66; mean ± SD_hearing_ = 37.15 ± 16.85; t_(42)_ = 0.74, p = .46), gender (deaf: 15 females/9 males; hearing: 15 females/5 males; Χ^2^_(1)_ = 0.10, p = .76), reasoning skills (the Block Design subtest of the Wechsler Abbreviated Scale of Intelligence, or WASI [Wechsler, 1999]; mean ± SD_deaf_ = 59.62 ± 8.69; mean ± SD_hearing_ = 57.47 ± 8.02 [n = 19], t_(41)_ = 0.83, p = .41), and visuospatial memory span (the computerised Corsi Block-tapping task [Kessels et al., 2000], implemented in the Psychology Experiment Builder Language software [PEBL; Mueller & Piper, 2014]; mean ± SD_deaf_ = 5.29 ± 0.75; mean ± SD_hearing_ = 5.40 ± 1.10 [n = 19], t_(41)_ = -0.36, p = .72).

**Table 1.** Language background of the deaf participants included in the current study.

| ID | Native language | Preferred language | Other languages |
| --- | --- | --- | --- |
| 1 | BSL | BSL | English |
| 2 | BSL | BSL | English |
| 3 | English, BSL | English | None |
| 4 | BSL | BSL | English |
| 5 | BSL | BSL | American Sign Language |
| 6 | English | English | BSL |
| 7 | English | English | None |
| 8 | Australian Sign Language | Australian Sign Language | BSL,<br>American Sign Language |
| 9 | English | English | BSL |
| 10 | English | English | BSL |
| 11 | BSL | BSL | English |
| 12 | Gesture / home sign | BSL | English |
| 13 | English | English | BSL |
| 14 | English | BSL | None |
| 15 | English | BSL | None |
| 16 | Gesture / home sign, BSL | BSL | English |
| 17 | English | English | None |
| 18 | English | BSL | None |
| 19 | English | BSL | None |
| 20 | English | English | None |
| 21 | English | BSL | None |
| 22 | English | English | None |
| 23 | English | English | BSL |
| 24 | English | BSL | South African Sign Language |

BSL, British Sign Language.

The ethics committee of the School of Psychology at the University of East Anglia and the Research and Development Department of the Norfolk and Norwich University Hospital approved the experimental procedure. All participants gave written informed consent and were compensated for their time. Participants who travelled to Norwich to take part in the study were reimbursed for their travel and accommodation expenses.

### Language assessment

We used grammaticality judgement tasks in English and BSL to measure the language skills of deaf participants and create a modality-independent measure of their general language proficiency. The design of the BSL Grammaticality Judgement Task (BSLGJT) is described in Cormier et al. (2012). The English version of the task (English Grammaticality Judgement Task; EGJT) was developed for this study by adapting stimuli sentences from Linebarger et al. (1983) to a computerised format. The tasks were implemented in PsychoPy (Peirce, 2007). Deaf participants completed both tasks if they knew both languages, or the English task only if they did not know BSL.

Accuracy measures (% of correct responses) from both tasks were used to create modality-independent language scores. Accuracy scores were z-transformed for each task separately, and the higher z-score of the two was chosen as a measure of general language proficiency for each participant (Figure 1). For example, a participant who had an accuracy of 53.93% and a z-score of -2.59 in the EGJT, and 78% and 0.07 respectively in the BSLGJT, would receive 0.07 as their normalised measure of their general language ability, reflecting proficiency in their stronger language (BSL). For four participants who did not know BSL, the EGJT score was used. Data from two participants were more than two SDs below the mean and were excluded from language-related analyses as outliers.

**Figure 1.**
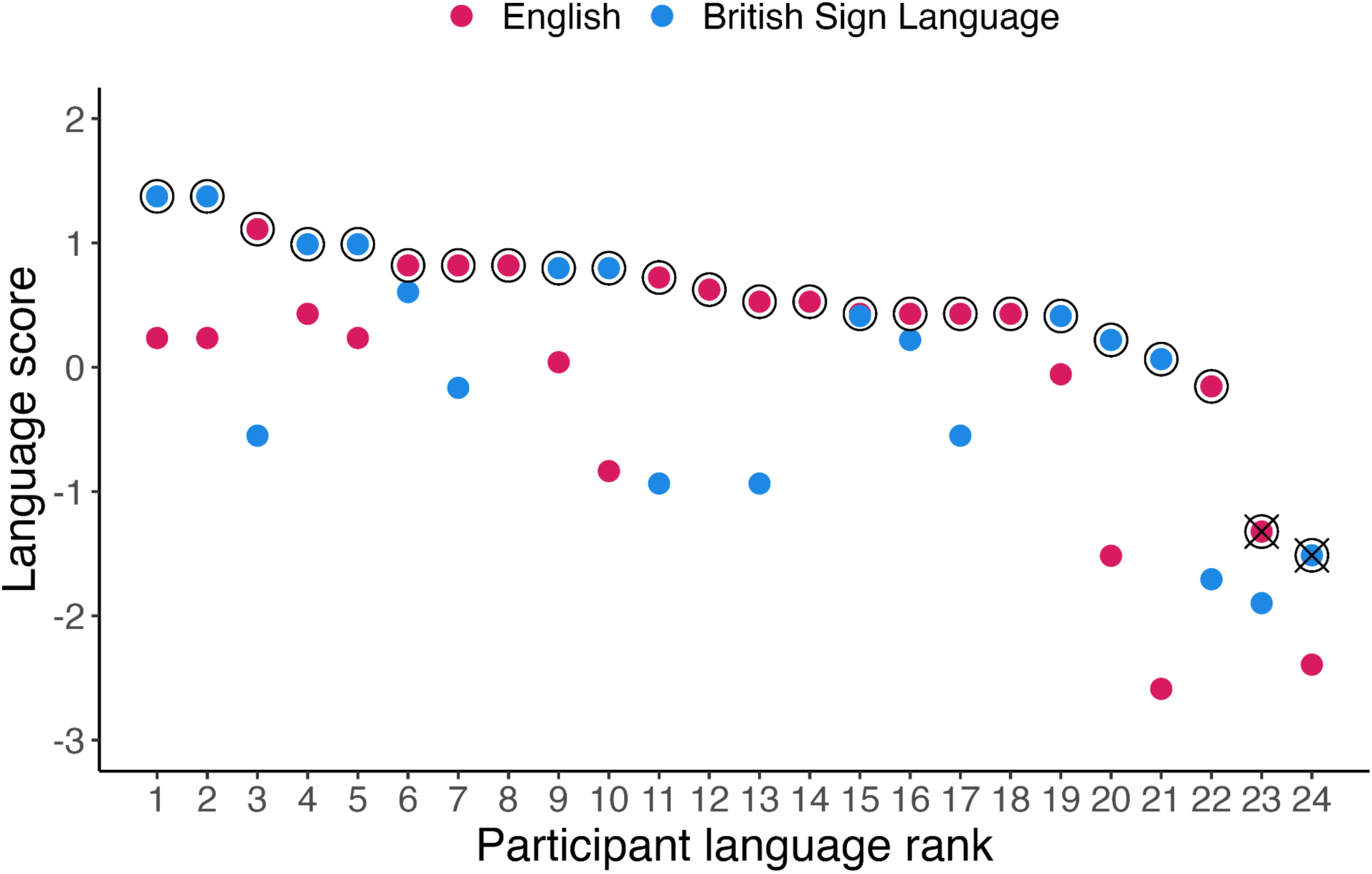
Modality-independent language scores of deaf participants. The higher language z-score, which was included into the modality-independent measure of language proficiency, is circled for each participant. Language data from participants with the lowest z-scores were excluded from the analyses as outliers and are marked with crossed circles.

### Experimental design and procedure

Each participant performed a scanning session, consisting of four experiments (switching, working memory, planning, and inhibition), a resting-state scan, and a behavioural session. The sessions took place on the same day or across two days. Data from these experiments have been previously reported in Manini et al. (2022). In this study, we included data only from the working memory and planning experiments. This is because our aim was to measure how responses and connectivity in the TP and TN networks changed as a function of the task participants were performing. To achieve this, we needed to compare two contrasting tasks with different executive demands. The working memory and planning experiments were selected because they included two distinct tasks: one that tapped the relevant EF (working memory, planning) and a control task, where executive demands were significantly lower (perceptual judgement [colour], counting beads). The switching and inhibition experiments did not meet the requirement for two contrasting tasks. In these experiments, participants performed the same task responding to identical visual features throughout, with conditions that varied only in the level of executive demands.

Instructions were given in either English or BSL, depending on the preference of each participant. Participants also received a written description of the tasks. Pre-scanner training ensured that participants were familiar with the procedure and understood the tasks. Participants were required to achieve an accuracy rate of at least 75% correct responses before scanning.

*Working memory.* The visuospatial working memory experiment was adopted from Fedorenko et al. (2011, 2013), with an added perceptual control (colour) condition (Figure 2). In both tasks, black squares were displayed for 1000 ms three times per trial at random locations on a 3 x 4 grid. A visual cue appeared before each trial to denote which task the participant had to perform (working memory or control [colour]). In the working memory condition, participants had to memorise the positions of the black squares and choose one of two alternative-response 3 x 4 grids (by pressing a corresponding left or right button). The grids were displayed for a maximum of 3750 ms or until a response button was pressed. In the colour task, participants were asked to memorise if a blue square appeared on any of the three grids throughout the trial and respond with a ‘yes’ or ‘no’ answer. The ‘yes’ and ‘no’ responses were presented on the left and right side of the screen and corresponded to the left and right buttons. The duration of the inter-trial interval was jittered between 2000 ms and 3500 ms. Each run of the task included 30 working memory trials and 30 control trials.

**Figure 2.**
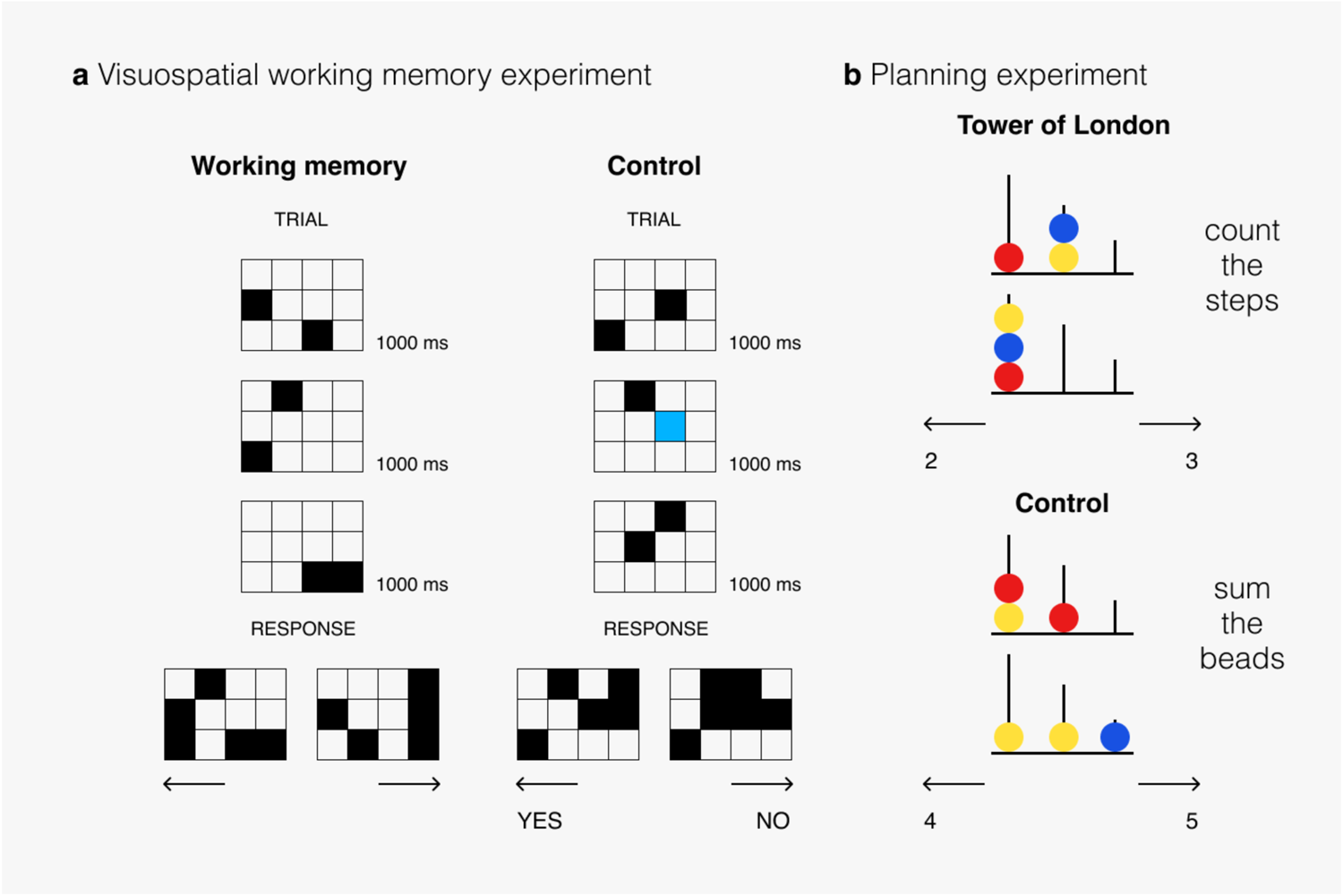
Schematic illustrations of the stimuli and design used in the working memory and planning experiments. The figure displays the types of tasks: working memory and control (colour), and planning and control (counting beads).

*Planning.* The planning task included a computerised version of the Tower of London task (Figure 2). The experimental paradigm was adopted from den Heuvel et al. (2003). For each trial, text at the top of the display screen indicated the task participants had to perform. In the planning task, participants looked at two configurations of three coloured beads on three vertical rods. They were asked to determine the minimum number of steps required to transform the upper configuration into the lower one (the target configuration). Participants followed two rules: (i) they could move only one bead at a time; (ii) they could not move a bead when there was another bead on top of it. The possible answers (the number of moves) were presented on the left and right sides of the screen and corresponded to left and right button responses. The level of complexity varied depending on the number of moves required (2, 3, 4, and 5). In the control task, participants counted how many blue and yellow beads were present on the screen during the trial. The inter-trial interval was jittered between 2000 ms and 3500 ms. The maximum trial duration was set to 3000 ms. There were 30 trials of each condition.

### Behavioural data analysis

The mean reaction time (RT) and percentage of correct responses were calculated for each participant and each task. To exclude outliers in RTs, the interquartile range was calculated as the difference between the upper and lower quartiles, and response values that were lower than 1.5 times the interquartile range from the lower quartile or greater than 1.5 times the interquartile range from the upper quartile, were removed.

Differences between the groups were tested with separate 2 x 2 ANOVAs for each experiment and type of response measure (accuracy, RT). The response accuracy rate (% of responses that were correct) and RT were used as dependent variables. Task was the within-subjects factor (working memory and colour in the working memory experiment, and planning and counting beads in the planning experiment), and group was the between-subjects factor (deaf, hearing).

To investigate the effect of language on behavioural measures, correlation coefficients (r) were calculated between modality-independent language scores in the deaf group and response values (accuracy, RT) in each task. The Bonferroni method was used to correct for multiple comparisons when appropriate. All statistical analyses were performed using JASP (JASP Team, 2023).

### fMRI data acquisition and analysis

#### Image acquisition

For each experiment, MRI structural and functional brain images were obtained in a GE Discovery MR750W 3T MRI whole-body scanner equipped with a 64-channel head coil. Functional data were acquired with a gradient-recalled echo-planar imaging (EPI) sequence (50 slices, in-plane resolution = 3 x 3 mm, slice thickness = 2 mm, distance factor = 50%, repetition time [TR] = 3000 ms, echo time [TE] = 50 ms, field of view [FOV] = 192 x 192 mm). Anatomical images were acquired using a T1-weighted scan (IR-FSPGR, inversion time = 400 ms, in-plane resolution = 1 x 1 mm, slice thickness = 1 mm) at the beginning of the session. Raw B0 field map data were acquired with a 2D multi-echo gradient-recalled echo sequence (TR = 700 ms, TE = 4.4 and 6.9 ms, flip angle = 50°, matrix size = 128 x 128, FOV = 240 x 240 mm, number of slices = 59, slice thickness = 2.5 mm, slice gap = 2.5 mm). Real and imaginary images were reconstructed for each TE to permit calculation of B0 field maps in units of Hz (Fessler et al., 2005; Funai et al., 2008; Jezzard & Balaban, 1995).

Task presentation and response logging were implemented using PsychoPy (Peirce, 2007) on a MacBook Pro (Retina, 15-inch; Mid 2015). The task was projected on a screen by an AVOTEC Silent Vision projector (Silent Vision, Avotec) at the end of the magnet’s bore. The projector screen was presented to participants through a mirror mounted on the head coil. Fibre-optic response pad boxes were used to record performance. Foam pads were used to reduce motion. All participants had ear protection. Deaf and hearing participants could communicate with the researchers in spoken or sign language (through a microphone or camera).

#### fMRI preprocessing

Functional volumes were preprocessed with SPM12 (Welcome Department of Imaging Neuroscience, London, UK), implemented in MATLAB 2018a (MathWorks, Inc.). Anatomical scans were bias-corrected and skull-stripped (see Manini et al., 2022 for details on preprocessing) and normalised to the standard Montreal Neurological Institute-Hospital (MNI) space. The deformation field from this step was used for the spatial normalisation of the functional scans. Susceptibility distortions in the echo-planar images were estimated using a field map that was co-registered to the blood oxygenation level-dependent (BOLD) reference (Fessler et al., 2005; Funai et al., 2008). Preprocessing steps included realignment (using the pre-calculated phase map), slice timing correction, co-registration, and smoothing (with a Gaussian kernel of 8 mm full-width at half-maximum). Functional scans were checked for excessive motion and artefacts using the Artifact Detection Tool (ART) within the CONN functional connectivity toolbox (Whitfield-Gabrieli & Nieto-Castanon, 2012) in SPM12.

#### Denoising

Prior to the connectivity analysis, functional data were denoised using the default CONN denoising pipeline, which includes detrending and a component-based noise correction method (CompCor; Behzadi et al., 2007). This approach removes subject-specific movement and physiological noise factors. White matter signals (5 CompCor components), cerebrospinal fluid signals (5 CompCor components), estimated subject-motion parameters (Friston et al., 1996), motion outlier scans detected with ART during scrubbing (Power et al., 2014), constant and first-order linear session effects, and constant task effects were regressed out during denoising. This was followed by band-pass filtering of the BOLD time series (0.008-0.09 Hz) (Whitfield-Gabrieli & Nieto-Castanon, 2012).

#### fMRI data analyses

During first-level analysis, a general linear model (GLM) encompassing the design and contrasts at the individual participant level was fitted for each experiment. The design events were modelled as boxcar functions and convolved with the canonical haemodynamic response function of SPM12. Button responses (separately for each condition and hand) and head motion parameters derived from the realignment of the images were included in the model as regressors of no interest.

*Working memory.* The model included information about the tasks (working memory, colour), onset times (set at the presentation of the first grid), and durations (fixed at 3500 ms, i.e., the duration of the presentation of the three grids [3000 ms] and the blank screen [500 ms] before the response window appears).

*Planning.* The model included information about conditions (planning, counting beads), onset times (set at the beginning of each trial), and durations (set to the trial-specific RT).

#### Regions of interest (ROIs)

ROIs for the TP and TN networks were adapted from a study by Fox et al. (2005), which identified these regions based on the analysis of resting-state functional data and replicated the findings across different rest conditions (visual fixation, eyes closed, and eyes opened). TP regions were centred in the frontal eye field (FEF) region of the precentral sulcus [left: -24 -12 61] [right: 28 -7 54], dorsolateral prefrontal cortex (DLPFC) [left: -40 39 26] [right: 38 41 22], inferior parietal lobule (IPL) [left: -42 -44 49] [right: 47 -37 52], and inferior precentral sulcus (IPS) [left: -23 -66 46] [right: 25 -58 52]. Default mode, or task-negative (TN), regions were defined in the retrosplenial cortex (RSC) [3 -51 8], posterior cingulate/precuneus (PCC) [-2 -36 37], superior frontal (SF) [left: -14 38 52] [right: 17 37 52], and lateral parietal cortex (LP) [left: -47 -67 36] [right: 53 -67 36]. ROIs were defined as spheres of 10 mm radius centred around the reported peak coordinates, using the MarsBaR toolbox for SPM (Brett et al., 2002; http://marsbar.sourceforge.net/). The mean contrast values were extracted for each participant and each ROI in the MarsBar toolbox from the relevant contrast images (working memory, colour; planning, counting beads; each relative to baseline) generated during the first-level analysis. The computed mean contrast values were averaged across ROIs from the TP and TN networks and used later in second-level analyses.

#### Functional connectivity analyses

Functional connectivity analyses were performed in CONN. ROI-to-ROI connectivity matrices were estimated for each task, characterising the connectivity between the 14 ROIs. The strength of connectivity between the BOLD time series of the ROIs in each task was defined from a weighted GLM (Nieto-Castanon, 2020) as Fisher z-transformed correlation coefficients.

#### Statistical analyses

Mean contrast values (averaged across regions in each network and participants in each group) were entered into 2 x 2 x 2 ANOVAs with within-subjects factors network (TP, TN), task (EF task [working memory or planning], control task [colour or counting beads]), and the between-subjects factor group (deaf, hearing).

For functional connectivity analyses, ROI-to-ROI functional connectivity correlation coefficients were averaged to get within-network (TP, TN) and between-network (TP-TN) connectivity measures and entered into 2 x 2 x 2 ANOVAs with within-subjects factors network (TP, TN, TP-TN), task (EF task [working memory or planning], control task [colour or counting beads]), and the between-subjects factor group (deaf, hearing). The Greenhouse-Geisser sphericity correction was applied when the assumption of sphericity was violated.

To investigate the effect of language on TP and TN networks, correlation coefficients (r) were calculated between modality-independent language scores in the deaf group and mean contrast values or connectivity values. Significance levels for correlation coefficients were adjusted for multiple comparisons according to the Bonferroni method.

## Results

### Behavioral data

Table 2 shows the accuracy (% of correct responses) and RT in the working memory and planning experiments.

**Table 2.** Accuracy and RT in the working memory and planning experiments.

| **Accuracy** |  |  |  |  |
| --- | --- | --- | --- | --- |
| Mean ± SD (% correct) | | | | |
| **Experiment** | **Group** |  |  |  |
|  | Deaf |  | Hearing |  |
| Working memory | **EF** | **Control** | **EF** | **Control** |
| | 84.01 ± 9.73 | 97.59 ± 3.22 | 82.31 ± 9.7 | 97.95 ± 4.0 |
| Planning | **EF** | **Control** | **EF** | **Control** |
| | 77 ± 10.00 | 93.88 ± 11.47 | 74.62 ± 11.85 | 91.52 ± 11.92 |
| **Reaction time** |  |  |  |  |
| Mean ± SD (seconds) | | | | |
| **Experiment** | **Group** |  |  |  |
|  | Deaf |  | Hearing |  |
| Working memory | **EF** | **Control** | **EF** | **Control** |
| | 1.64 ± 0.39 | 0.88 ± 0.29 | 1.48 ± 0.4 | 0.68 ± 0.13 |
| Planning | **EF** | **Control** | **EF** | **Control** |
| | 7.91 ± 1.43 | 3.95 ± 1.52 | 7.07 ± 1.92 | 2.7 ± 0.47 |

\*Mean RT across all correct trials.
RT, reaction time.

Accuracy did not differ between the groups in either experiment. The main effect of task was significant in both experiments (working memory: F_(1,40)_ = 87.12, p < .001, planning: F_(1,37)_ = 42.98, p < .001).

A significant main effect of task was also observed for RT in both experiments (working memory: F_(1,40)_ = 189.74, p < .001; planning: F_(1,37)_ = 232.95, p < .001). There was also a significant main effect of group for RT (working memory: F_(1,40)_ = 4.57, p = .04; planning: F_(1,37)_ = 7.9, p = .008), but no significant interactions involving group. Post-hoc two-tailed t-tests revealed that the deaf group had overall significantly slower RT in both experiments: working memory (mean ± SD_deaf_ = 1.26 ± 0.29, mean ± SD_hearing_ = 1.08 ± 0.25, t_(40)_ = 2.14, p = .04l) and planning (mean ± SD_deaf_ = 5.93 ± 1.21, mean ± SD_hearing_ = 4.89 ± 1.11, t_(37)_ = 2.81, p = .008).

Language scores did not significantly correlate with any of the EF measures. However, they significantly correlated with RT (r = -.60, p = .008; Bonferroni-corrected for four comparisons, p < .01) in the control (counting beads) condition of the planning experiment, suggesting faster response times in the control task in deaf participants with higher language scores (Figure S1). Correlation between language scores and accuracy in the same condition did not survive the Bonferroni correction (p = .016, Bonferroni-corrected p-value threshold = .01) (see Figure S1 in the supplementary materials for scatterplots of the relationships between language scores and behavioural measures).

### Working memory

*Univariate fMRI analysis.* First, we investigated between-group differences in activation patterns in the task-dependent regions. We ran an ANOVA on contrast values with the following factors: network (TP, TN), task (working memory, colour), and group (hearing, deaf). The main effects of network (F_(1,40)_ = 29.38, p < .001) and task (F_(1,40)_ = 5.44, p = .03) were significant. The interactions between network and task (F_(1,40)_ = 108.60, p < .001) and network, task, and group (F_(1,40)_ = 6.36, p = .02) were also significant. As expected, the mean contrast values were significantly higher during the cognitively demanding working memory (WM) task, compared to the control colour task, in the TP regions (mean_WM_ ± SD = 1.85 ± 2.54; mean_control_ ± SD = 0.07 ± 2.12,; t_(41)_ = 8.14, p < .001, while the TN regions showed a larger decrease in activations during the working memory task (mean_WM_ ± SD = -1.90 ± 2.10; mean_control_ ± SD = -0.98 ± 1.73; t_(41)_ = -4.41, p < .001).

To explore the interaction involving the factor group, we conducted separate 2 (task: working memory, control) x 2 (group: hearing, deaf) ANOVAs for each network. A significant main effect of task was found in both networks: the TP network (F_(1,40)_ = 51.48, p < .001) and the TN network (F_(1,40)_ = 27.26, p < .001). There was a significant task x group interaction in the analysis of the TN network (F_(1,40)_ = 9.07, p = .004). The interaction was driven by a significant difference between the working memory and colour tasks in the hearing group (t_(19)_ = -5.56, p = <.001, post-hoc two-tailed t-tests; Bonferroni-corrected for six comparisons, p < .008) (Figure 3), suggesting deactivation of the TN regions during the control (colour) task in the hearing group, but not in the deaf group.

*Language effects.* The main aim of the study was to investigate what the variability of language skills in the deaf group could reveal about the function and connectivity of the task-related regions during cognitive tasks. To examine whether language scores are associated with changes in the BOLD signal during the working memory experiment in the deaf group, we correlated activation in each task (working memory, colour) in each network (TP, TN) with language proficiency scores. Only activations during the working memory task in the TP network showed a significant positive correlation with language proficiency scores, with the Bonferroni correction for four comparisons applied (r = .56, p = .009; Bonferroni-corrected p-value threshold: p = .01) (Figure 4; see Figure S2 in the supplementary materials for associations that were not statistically significant).

**Figure 3.**
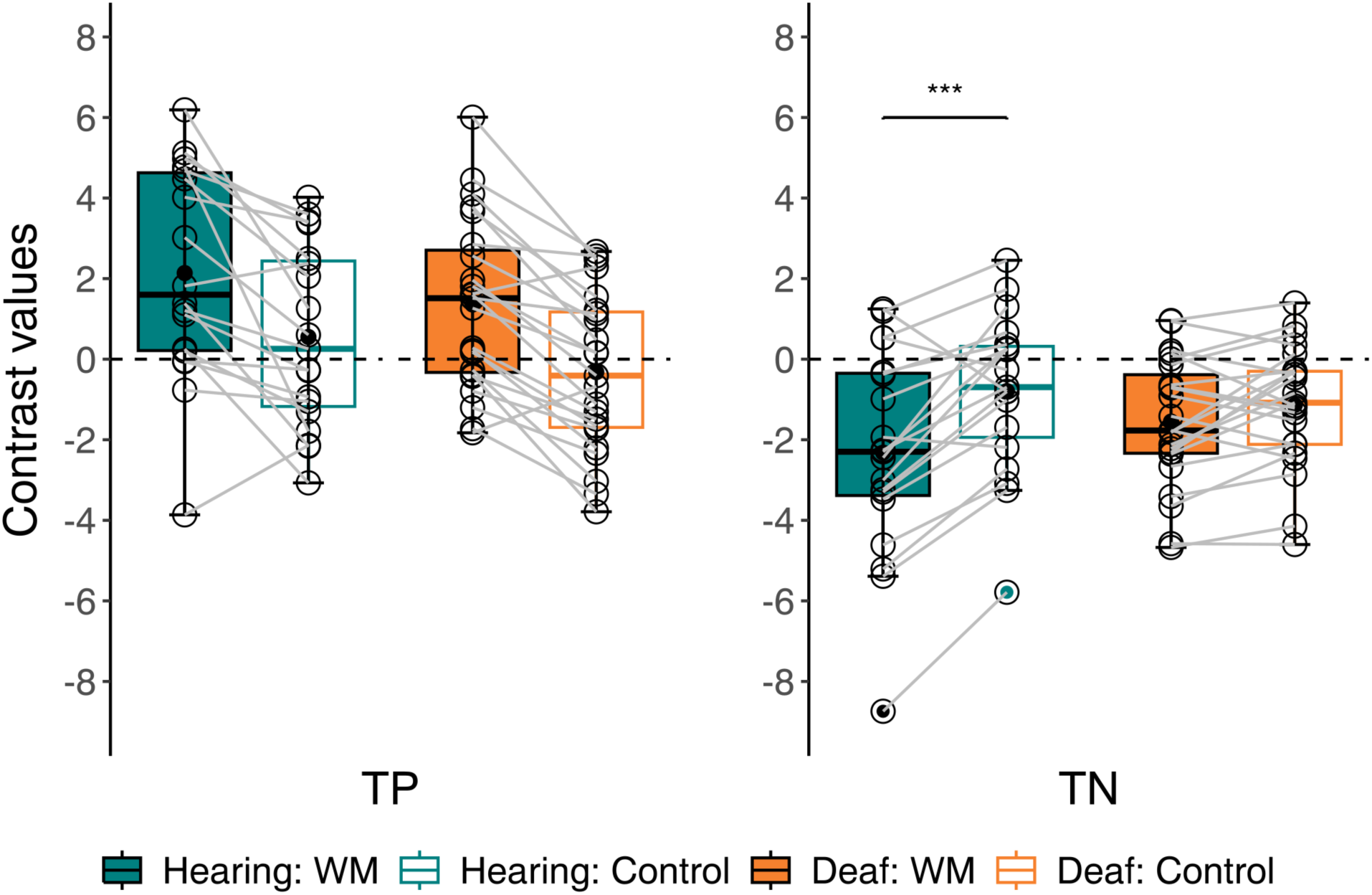
Comparison of working memory activations between hearing and deaf participants. Boxplots show mean contrast values for the working memory and control (colour) tasks, averaged across regions from the TP and TN networks in the deaf and hearing groups. The task x group interaction in the TN regions was significant (F_(1,40)_ = 9.07, p = .004). Post-hoc t-tests revealed a significant difference between the working memory and control tasks in the hearing group only (t_(19)_ = -5.56, p < .001). WM, working memory; TP, task-positive; TN, task-negative.

**Figure 4.**
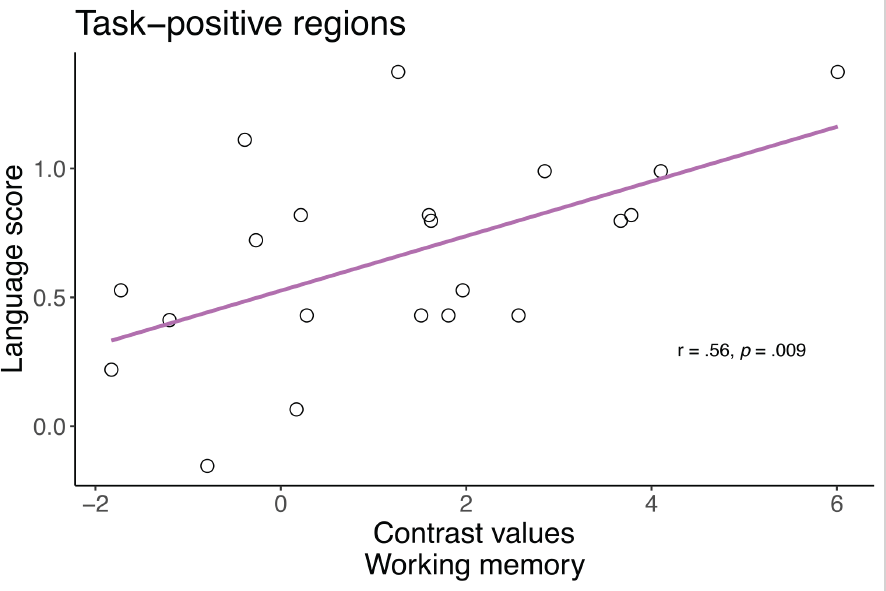
Correlation between language scores and working memory activity in the task-positive network. Scatterplot of language proficiency scores plotted against contrast values from the TP regions during the working memory task in the deaf group. The positive correlation between language scores and contrast values was statistically significant (r = .56, p = .009).

*Functional connectivity.* In the analysis of differences in functional connectivity between the groups during the working memory experiment, the main effects of network (F_(2,80)_ = 143.53, p < .001) and group (F_(1,40)_ = 8.44, p = .006) were significant. Post-hoc two-tailed t-test comparisons indicated that the deaf group had significantly higher overall connectivity across tasks and networks (t_(40)_ = 2.91, p = .006; mean ± SD_deaf_ = 0.17 ± 0.05; mean ± SD_hearing_ = 0.13 ± 0.04) (Figure 5).

*Language effects on functional connectivity.* To explore the effect of differences in language proficiency levels on functional connectivity in the deaf group, we calculated correlation coefficients between language scores and connectivity in each task and each network.

There was a significant correlation between language scores and connectivity only in the TP network during the working memory task (r = .64, p = .002; Bonferroni-corrected p-value threshold: p = .01) (Figure 6). All other correlations were not significant (see Figure S3 in the supplementary materials).

**Figure 5.**
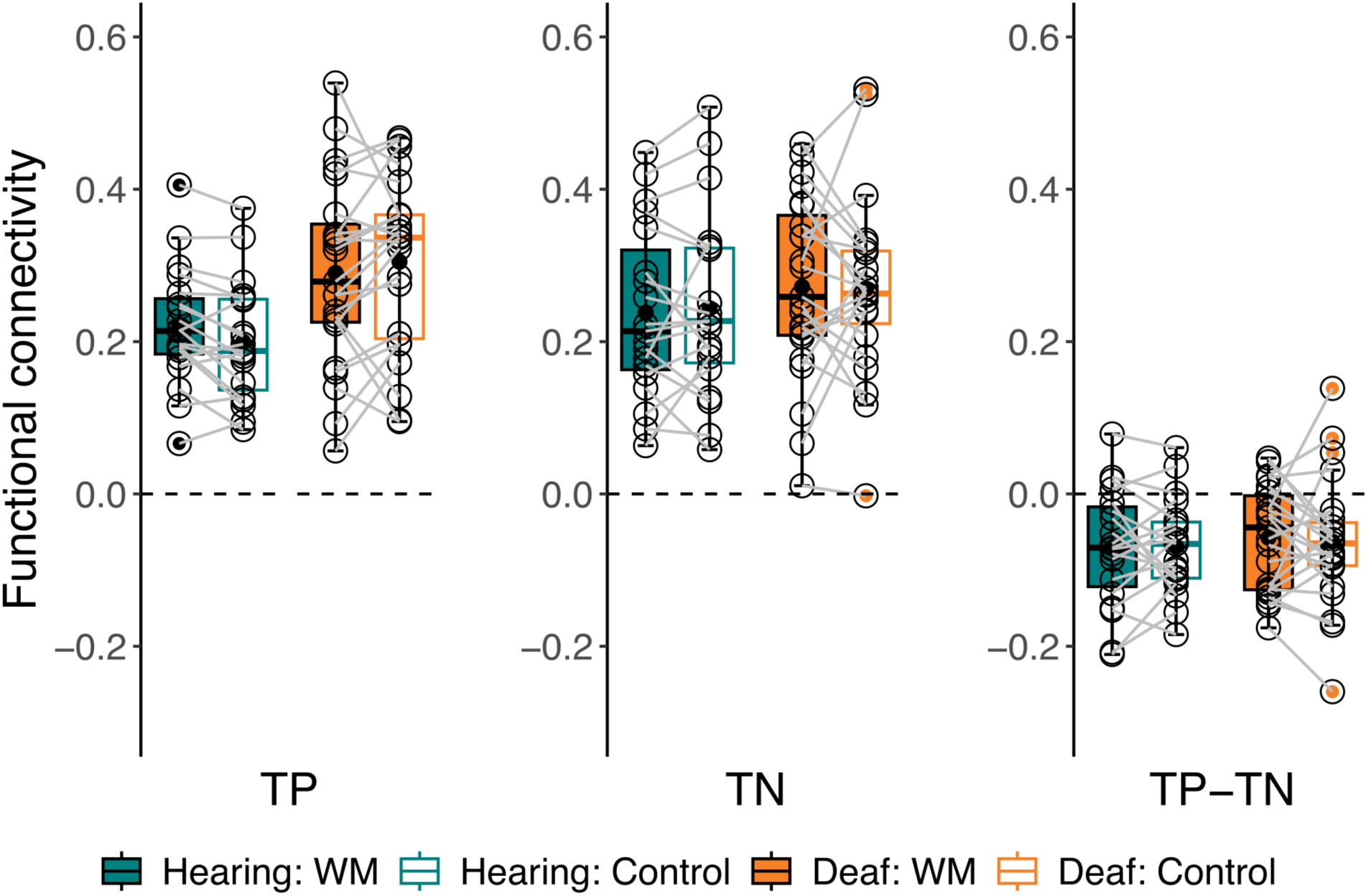
Functional connectivity in the working memory experiment in deaf and hearing participants. Boxplots show mean task-related functional connectivity (Fisher z-transformed r-values) for the TP, TN, and TP-TN connections in each task of the working memory experiment (working memory, colour), averaged in each group. Repeated-measures ANOVA revealed a main effect of group (F_(1,40)_ = 8.44, p = .006), and a Bonferroni-corrected post-hoc t-test confirmed that the overall connectivity was significantly higher in the deaf group (t_(40)_ = 2.91, p = .006). WM, working memory; TP, task-positive; TN, task-negative.

**Figure 6.**
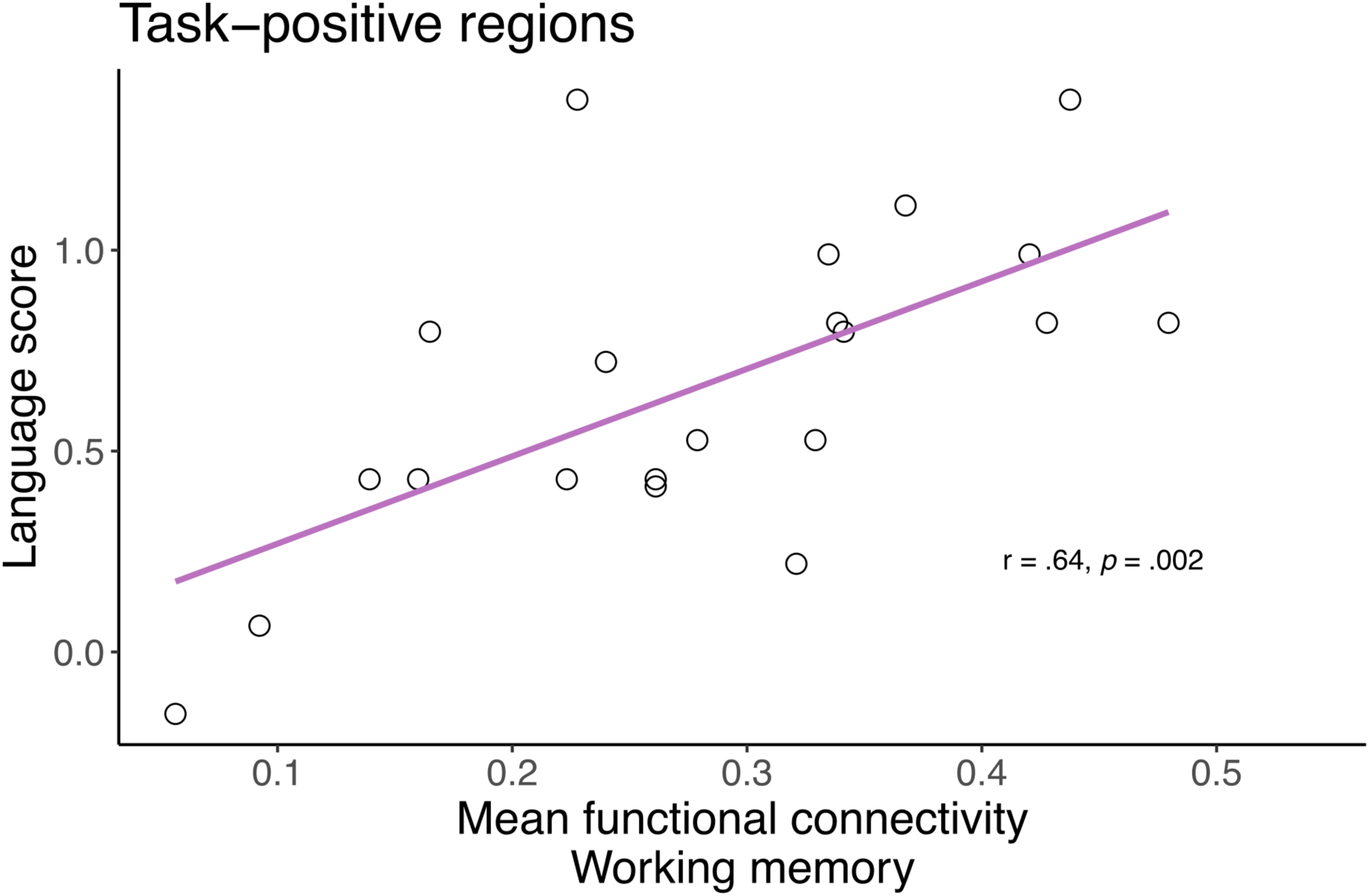
Correlation between language scores and functional connectivity during working memory in the task-positive regions. Scatterplot of language proficiency scores against functional connectivity values for the working memory task in the TP regions in the deaf group. The positive correlation between language scores and functional connectivity in the working memory task in the TP regions was statistically significant (r = .64, p = .002).

### Planning

*Univariate fMRI analysis.* The ANOVA did not reveal any significant effects or interactions with group. The main effects of task (F_(1,37)_ = 30.34, p < .001) and network (F_(1,37)_ = 210.46, p <.001) were significant (Figure 7).

**Figure 7.**
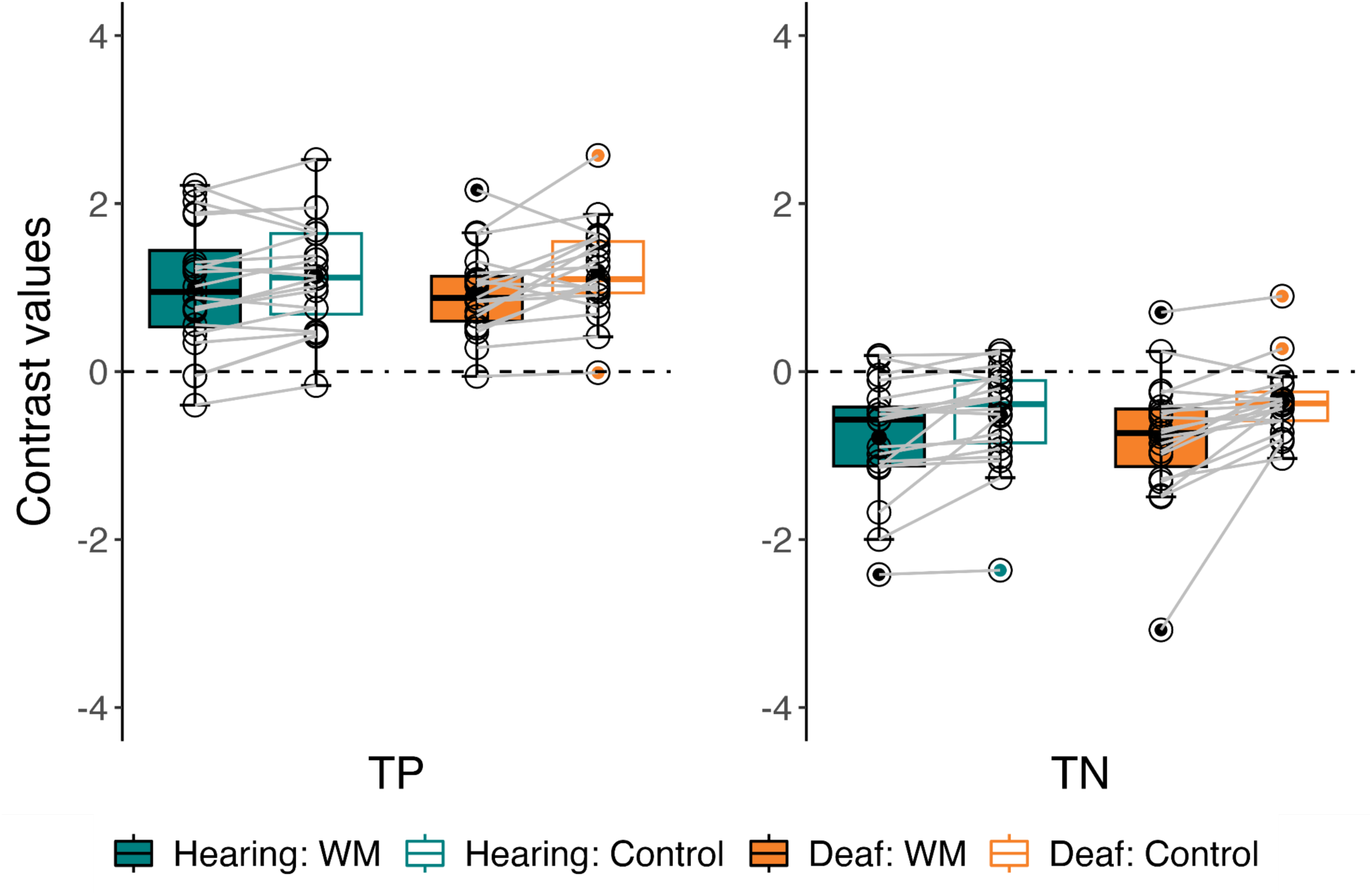
Activations in the planning experiment in the deaf and hearing groups. Boxplots display mean contrast values for the planning and control (counting beads) tasks, averaged across regions within the TP and TN networks for deaf and hearing participants. TP, task-positive; TN, task-negative.

*Language effects.* Correlations between language scores and the contrast values in each network and task combination were not significant (Figure S4).

*Functional connectivity.* The only significant effect in the connectivity analysis of the planning experiment was the effect of network (F_(2,70)_ = 150.55, p < .001) (Figure 8).

**Figure 8.**
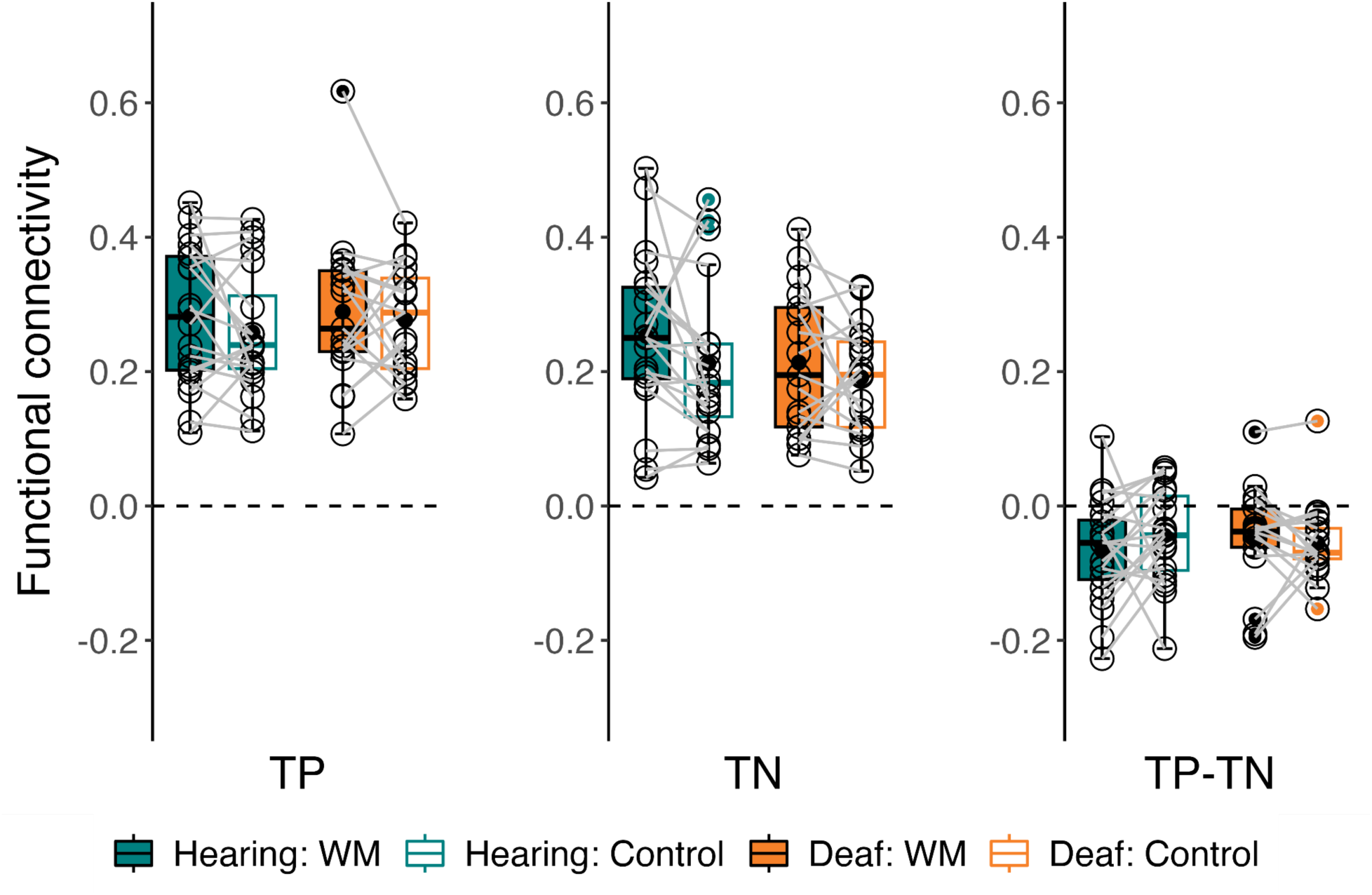
Functional connectivity in the planning experiment in hearing and deaf participants. Boxplots show mean task-related functional connectivity (Fisher z-transformed r-values) for the TP, TN, and TP-TN connections in each task of the planning experiment (planning, counting beads), averaged for each group. TP, task-positive; TN, task-negative.

*Language effects on functional connectivity.* Correlations between language scores and functional connectivity in each network and task combination were not significant in the planning experiment (Figure S5).

## Discussion

In this study, we examined the impact of differences in language proficiency on brain networks involved in cognitive task performance in congenitally and early deaf participants. Prior research has demonstrated the effects of sensory experience on executive processing in the brain (Cardin et al., 2018; Ding et al., 2015; Manini et al., 2022) and on the functional connectivity of networks supporting cognition (Andin & Holmer, 2022; Bonna et al., 2021; Cardin et al., 2023; Malaia et al., 2014). Our findings provide unique insights into how activity and functional connectivity in task-related regions during higher-order cognition can be modulated by language experience. Variability in early language experience in deaf individuals leads to substantial differences in language proficiency (Cormier et al., 2012; Mayberry & Eichen, 1991; Mayberry & Fischer, 1989), which, in turn, influences cognitive functioning.

We did not observe a significant correlation between language proficiency scores and performance in the working memory or planning tasks in our sample. However, differences in modality-independent language proficiency were positively associated with the recruitment and functional connectivity of brain regions involved in higher-order cognition during working memory. When participants perform a visuospatial working memory task, they engage a set of frontoparietal areas, including the frontal eye field, dorsolateral prefrontal cortex, inferior parietal lobule, and inferior parietal sulcus (Belger et al., 1998; Curtis, 2006; Niendam et al., 2012; Postle et al., 2000; Rowe et al., 2000). Higher activity in frontal and parietal regions is associated with working memory training, capacity, and performance (Klingberg et al., 2002; McNab & Klingberg, 2008; Ogawa et al., 2014; Olesen et al., 2004; Pessoa et al., 2002). Similarly, increased frontoparietal connectivity is a neural marker of working memory training and more accurate task performance (Shen, 2015; Thompson et al., 2016). Our results suggest increased efficiency in cognitive processing in deaf individuals with higher language skills, irrespective of the sensory modality of their stronger language. Additionally, our findings demonstrate that examining neural activity can unveil relationships between specific aspects of cognitive processing and language, even when these relationships are not evident behaviourally.

These findings are in line with theories of cognitive development that suggest causal influences of language and social interaction on cognition (Luria, 1979; Luria & Yudovich, 1971; Vygotsky, 1962; Zelazo et al., 2003). Language, as a rule-based system, has been proposed to be essential for formulating, integrating, and maintaining higher-order action-oriented rules (Zelazo, 2015; Zelazo et al., 2003). Evidence indicates that linguistic strategies can act as an internal self-cueing device and support understanding, retrieving, and activating representations of task operations (Emerson & Miyake, 2003; Kray et al., 2008). Complex task rules are encoded internally through inner speech and are held in working memory, allowing flexibility and control in various goal-directed contexts (Zelazo, 2015). The cognitive complexity and control theory suggests that children have to acquire and master the linguistic means of representing rule structures to be able to successfully perform sophisticated cognitive tasks (Zelazo et al., 1996). Furthermore, language allows children to evaluate and reflect on their actions and mental processes, leading to improvements in their cognitive control abilities (Lyons & Zelazo, 2011). In conclusion, developmental models propose that language can support the emergence and early advances in EF skills (Marcovitch & Zelazo, 2009). In our study, we demonstrate a link between language proficiency, shaped by early developmental experiences of deaf people, and task-associated brain processing, providing neuroimaging support for theories that emphasise the role of language in cognition.

The working memory task used in our study required manipulation and maintenance of visuospatial information. Research suggests that spatial cognition is closely linked to the development of spatial language. For example, in deaf signers of an emerging sign language in Nicaragua, consistent linguistic marking of spatial information was correlated with performance in nonverbal spatial tasks (Pyers et al., 2010). Similarly, deaf children who had not been exposed to a conventionalised language rarely produced gestures that coded spatial relations and showed at-chance performance during a spatial mapping task, while hearing children could perform the task successfully (Gentner et al., 2013). In hearing children, the amount of spatial talk with parents predicts their use of spatial language, which is associated with better performance in spatial tasks at a later age (Pruden et al., 2011). This evidence suggests a developmental relationship between parent spatial language input, children’s spatial language use, and spatial reasoning abilities.

Spatial language can provide a framework for formulating conceptual spatial representations and enable spatial reasoning (Gentner & Loewenstein, 2002; Loewenstein & Gentner, 2005). More broadly, language may act as a ‘cognitive toolkit’, supporting relational reasoning and problem-solving (Gentner, 2016). Models of cognitive development and empirical evidence suggest that language can support and facilitate cognitive function either as a means of representing information about task rules or as a tool for abstract representation and reasoning. In our study, we describe the association between differences in language proficiency and visuospatial cognition in the brain. We do not disentangle the effects of particular language components on brain function, and the grammaticality judgement tasks used in our study did not specifically target spatial language in either modality. Marschark et al. (2015) previously reported a link between language proficiency in the preferred modality and visuospatial cognitive abilities in deaf children. Overall, our findings and evidence from other studies on deafness may point to a more general effect of higher language proficiency across different sentence types and structures on cognitive function. It is also possible that this advantage may be related to syntactic processing. The acquisition of complex syntactic structures, particularly of the ‘if-then’ conditional type, has been proposed as a mechanism that facilitates EF in children (Zelazo et al., 2003).

‘If-then’ reasoning is involved in planning and problem-solving, which are assessed by the Tower of London task used in our study. Research has demonstrated that planning performance can be affected by articulatory suppression (Lidstone et al., 2010, 2012; Wallace et al., 2017). Given the theoretical and empirical support for the associations between language and planning, we expected that cognitive processing in task-related regions during the execution of the Tower of London task would be linked to differences in language proficiency. However, we did not find a significant relationship between these measures in our sample of deaf participants. Considering that the use of private speech in planning has been suggested to become internalised in middle childhood (Lidstone et al., 2010), it may be more difficult to uncover these effects in adults. Studies with multiple cognitive control tasks and measures of specific aspects of language processing in deaf individuals of different ages are needed to describe the interplay between language and cognition throughout the lifespan. Employing a wide battery of tasks could help identify the specific computational processes shared by language and nonverbal cognition, providing a more comprehensive picture of their interactions.

Our study also compared neural activations and functional connectivity during task execution between the deaf and hearing participants. Both groups showed similar activation patterns in the TP and TN (or DMN) regions during working memory and planning tasks. However, while the hearing participants deactivated the regions of the DMN significantly more during the working memory task compared to the control task, the deaf group did not show the same difference between tasks in this network.

In the general population, DMN suppression is often described as relevant for goal-direction cognition, potentially reflecting the suspension of functions of this network that are not relevant for task execution, such as mind-wandering (Anticevic et al., 2012; Gusnard & Raichle, 2001). Both increased activation of task-specific areas and decreased deactivation of the task-negative/DMN regions have been linked to performance in an attentional task, suggesting a possible engagement of different neural circuits, depending on the cognitive strategy employed (Lawrence et al., 2003). The decreased deactivation in the DMN may specifically support the perceptual control task processing in hearing participants, who may shift to a different cognitive strategy for the working memory task, while deaf participants may employ more consistent neural circuits and strategies across the experiment.

Functional connectivity was overall higher in the deaf group across tasks and networks. The DMN consistently shows differences in resting-state functional connectivity profiles and network reorganisation between deaf and hearing individuals (Andin & Holmer, 2022; Bonna et al., 2021; Cardin et al., 2023; Dell Ducas et al., 2021; Kumar et al., 2021), with stronger functional connectivity in deaf participants (Bonna et al., 2021; Dell Ducas et al., 2021; Malaia et al., 2014) and differences in modular organisation (Bonna et al., 2021).

It remains unclear whether these effects are due to differences in language or sensory experience between deaf and hearing individuals. Although we did not show a direct relationship between activity and connectivity of the DMN and language proficiency scores, it is still possible that the differential patterns of recruitment between the groups in the task-negative regions and the overall higher connectivity in the working memory experiment in the deaf group are related to language skills. This idea is supported by evidence from studies comparing deaf native, early or highly proficient signers and hearing speakers, which have not found changes in DMN recruitment during working memory tasks (Andin et al., 2021; Cardin et al., 2018; Ding et al., 2015). These findings suggest that when language acquisition is controlled across groups, the sensory experience of deafness alone does not impact the DMN. In this study, we may not be capturing language-related effects in the DMN due to the use of a language task tapping specifically into grammaticality judgements. This could be disambiguated in future studies evaluating other aspects of language processing and their effects on the DMN.

## Conclusions

In this study, we investigated the influence of language and sensory experiences on cognitive processing in the brain. We demonstrate that language proficiency, independently of language modality, shapes both the function and connectivity of task-related regions involved in cognitive processing. Successful language acquisition, leading to higher language competency later in life, may enable the use of more efficient computational mechanisms for solving cognitive tasks, such as working memory. Our findings suggest that the acquisition of a complex linguistic system can support nonverbal cognition in the brain. This can be due to the involvement of specific linguistic-based structures that facilitate the formulation and maintenance of complex rules required in higher-order cognitive tasks. Our results may reflect dynamic interactions or shared neural resources between specific aspects of language processing and cognition during development.

There are other early environmental factors that can significantly impact the formation of cognitive skills. Socioeconomic indicators, such as family income, have been linked to cognitive performance, as well as cortical brain volume and thickness in children (Tomasi & Volkow, 2021). In deaf children, early social interactions and development of intersubjectivity may be disrupted due to environmental factors, which in turn has implications for language and EF development (Morgan et al., 2021). Consistent with the mutualism theory, skills in one cognitive domain can drive the development of skills in the other, and vice versa (Griffiths et al., 2022; Kievit et al., 2017). As our study shows, neuroimaging can provide valuable insights into the role of early environmental factors in cognitive processing, which is not always possible with behavioural measures alone. Further neuroimaging research involving deaf participants from diverse language backgrounds and across different age groups can provide a deeper understanding of the complex interactions between language, cognition, and sensory experience throughout development and across the lifespan.

## Acknowledgments

The authors thank all participants who took part in this study.

## Footnotes

1 One deaf participant listed Australian Sign Language as their native language (Table 1). British, Australian, and New Zealand Sign Language have been referred to as a group (BANZSL) (Schembri et al., 2010): they are historically related (Johnston & Schembri, 2007), have a high degree of lexical similarity (McKee & Kennedy, 2000) and minor or no grammatical differences (see Schembri et al., 2010).

