## Supplementary materials for "The impact of language proficiency on task-dependent neural activity and functional connectivity: Insights from deafness"

**Figure S1. Language proficiency scores and behavioural measures (accuracy, RT) in the deaf group.** Scatterplots of the language proficiency scores against behavioural measures (accuracy and RT) in the working memory and planning experiments. The positive correlation between RT in the control task of the planning experiment and the language proficiency scores was significant ($p = .008$, Bonferroni-corrected p-value threshold: $p = .01$).

RT, reaction time.

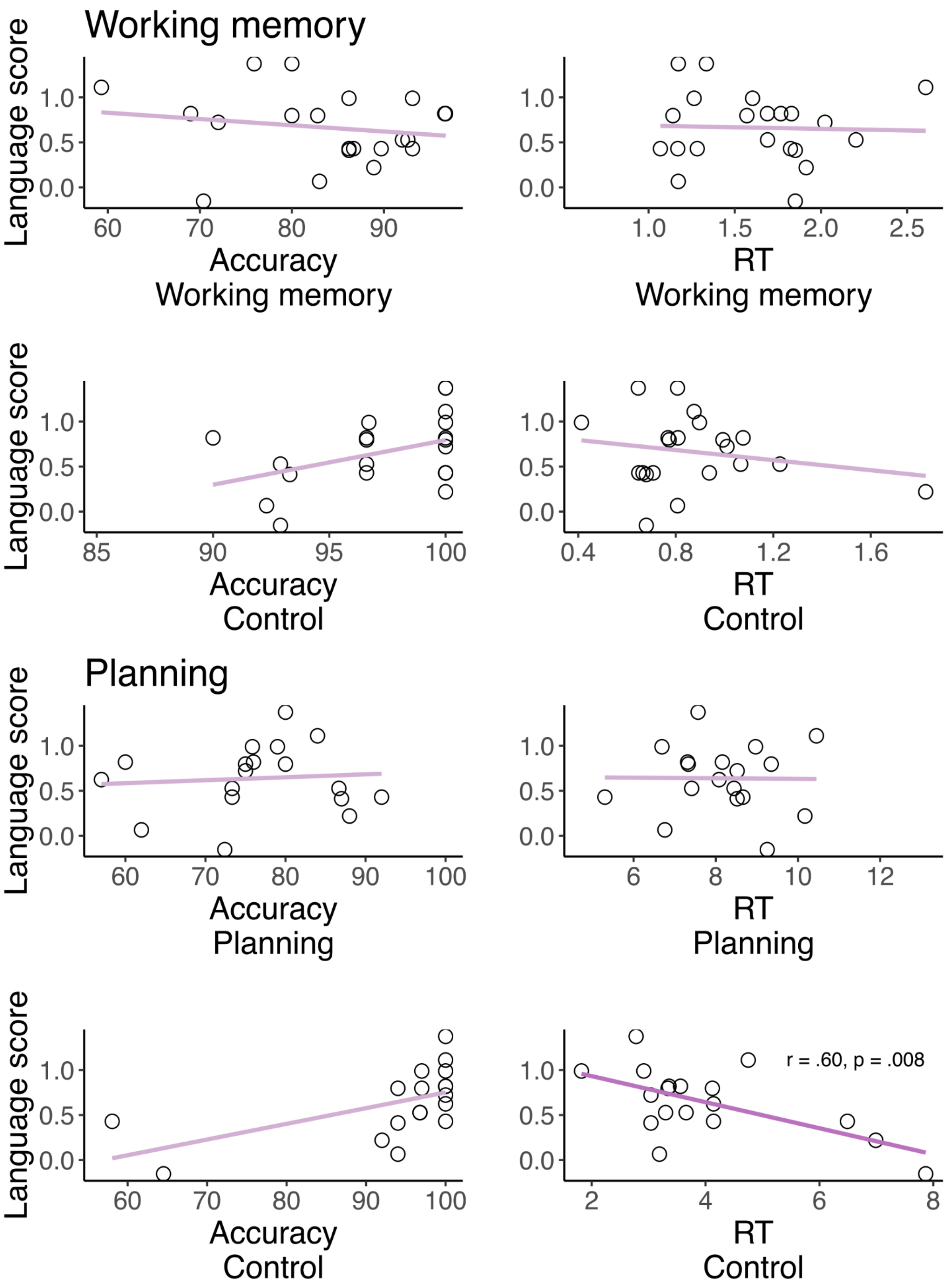


Note. After removing two outliers from the data on language scores and accuracy in the control task of the planning experiment (see the two leftmost points in the plot), the correlation coefficient changes to r = .62, p = .01; Bonferroni-corrected p-value threshold: p = .01).

**Figure S2. Language proficiency scores and activations in the working memory experiment in the deaf group.** Scatterplots of language proficiency scores against contrast values in the TP and TN regions during the working memory experiment. The correlations were not significant (Bonferroni-corrected p-value threshold: p = .01).
TP, task-positive; TN, task-negative.

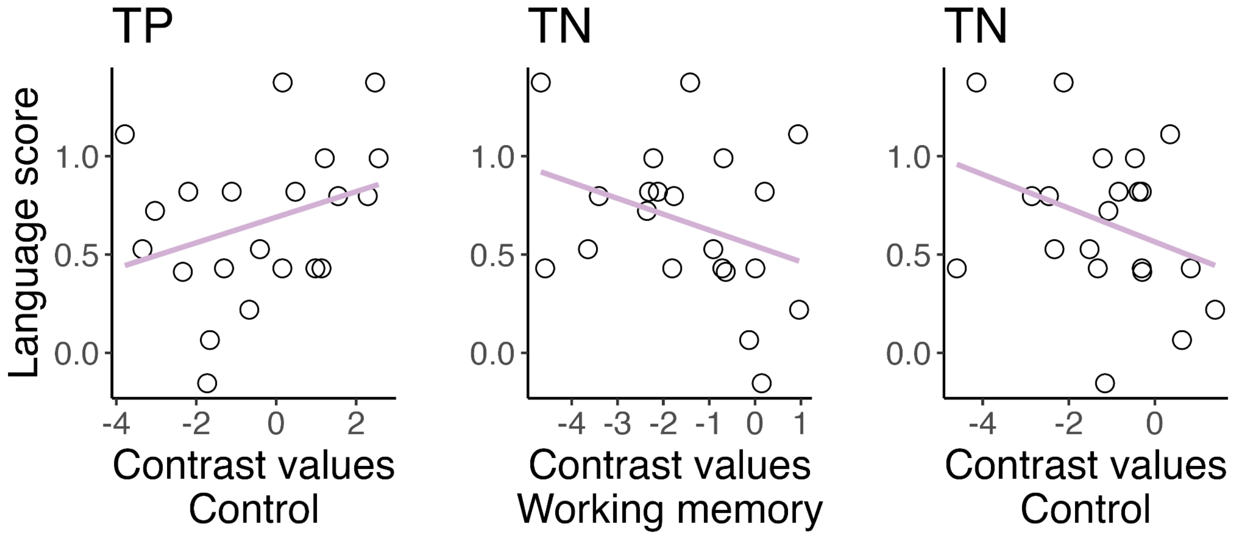

**Figure S3. Language proficiency scores and functional connectivity in the working memory experiment.** Scatterplots of language proficiency scores against within- and between-network functional connectivity in the TP and TN regions during the working memory experiment in the deaf group. The correlations were not significant (Bonferroni-corrected p-value threshold: $p = .01$).

TP, task-positive; TN, task-negative.

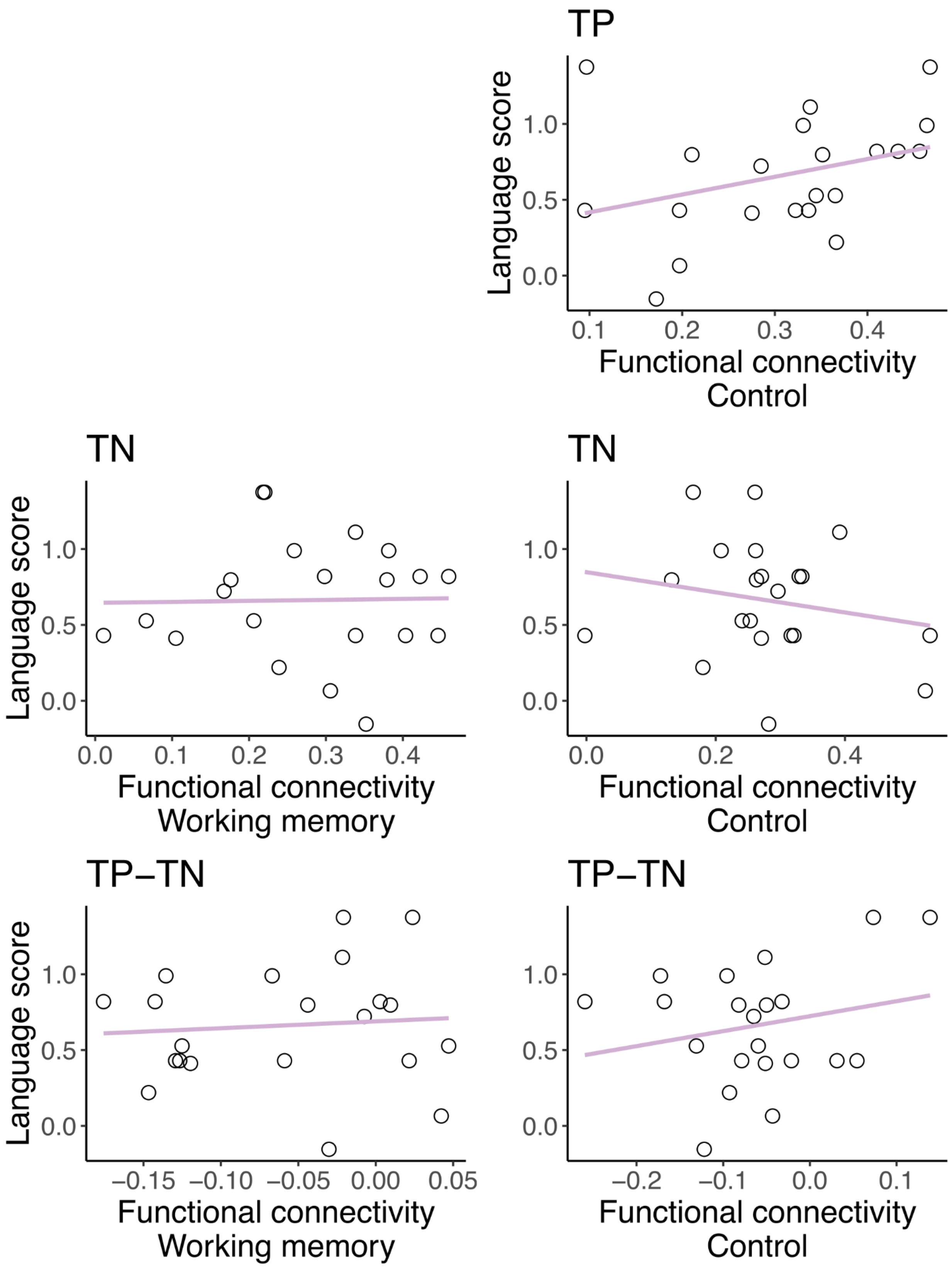

TP
Language score
1.0
0.5
0.0
0.1
0.2
0.3
0.4
Functional connectivity
Control
TN
Language score
1.0
0.5
0.0
0.0
0.1
0.2
0.3
0.4
Functional connectivity
Working memory
TN
1.0
0.5
0.0
0.0
0.2
0.4
Functional connectivity
Control
TP−TN
Language score
1.0
0.5
0.0
−0.15
−0.10
−0.05
0.00
0.05
Functional connectivity
Working memory
TP−TN
1.0
0.5
0.0
−0.2
−0.1
0.0
0.1
Functional connectivity
Control

**Figure S4. Language proficiency scores and activations in the planning experiment in the deaf group.** Scatterplots of language proficiency scores against contrast values in the TP and TN regions during the planning experiment. None of the corrections were significant (Bonferroni-corrected p-value threshold: p = .01).
TP, task-positive; TN, task-negative.

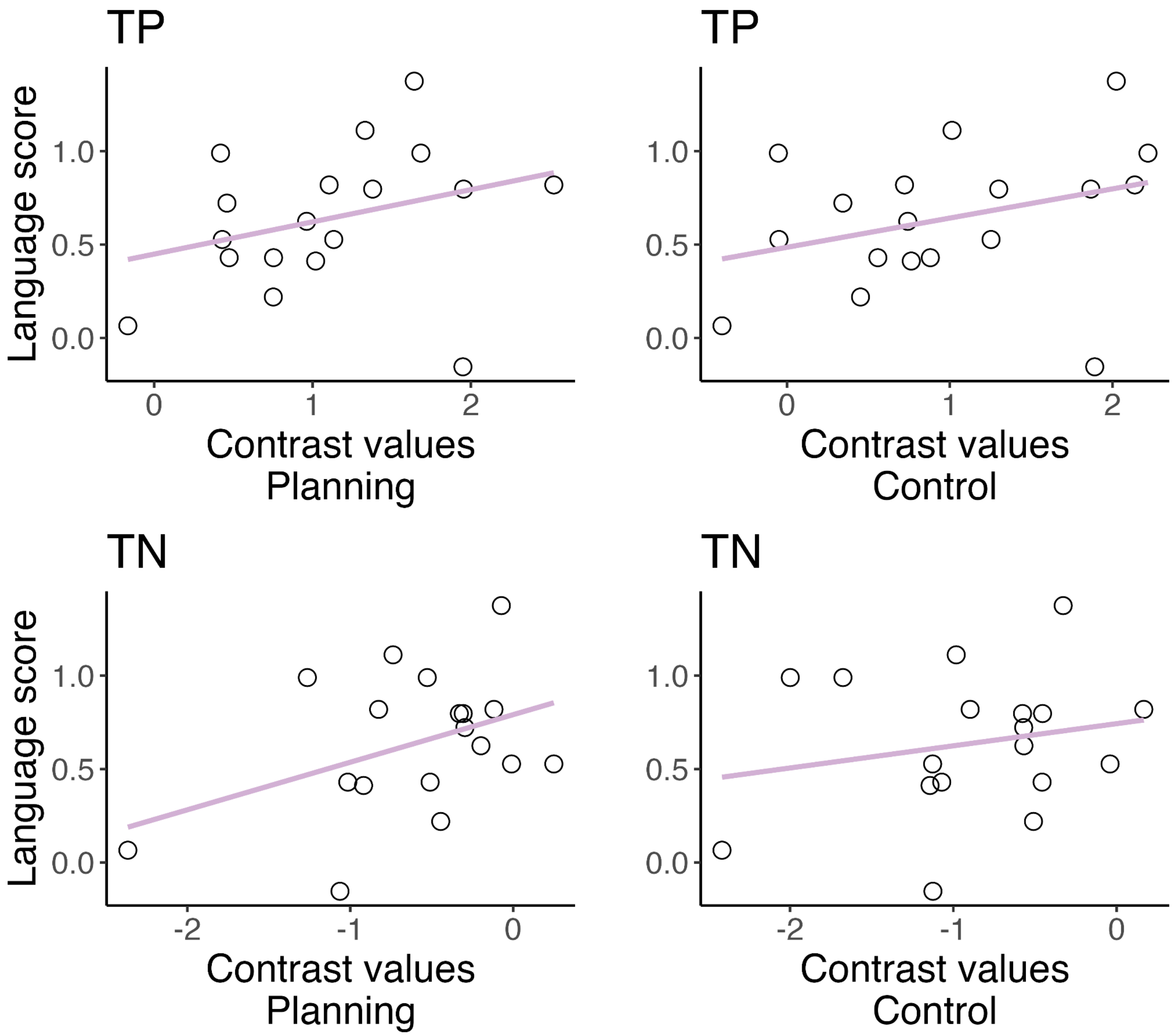

**Figure S5. Language proficiency scores and functional connectivity in the planning experiment.** Scatterplots of language scores against within- and between-network functional connectivity in the TP and TN regions during the planning experiment in the deaf group. None of the correlations were significant (Bonferroni-corrected p-value threshold: $p = .008$).

TP, task-positive; TN, task-negative.

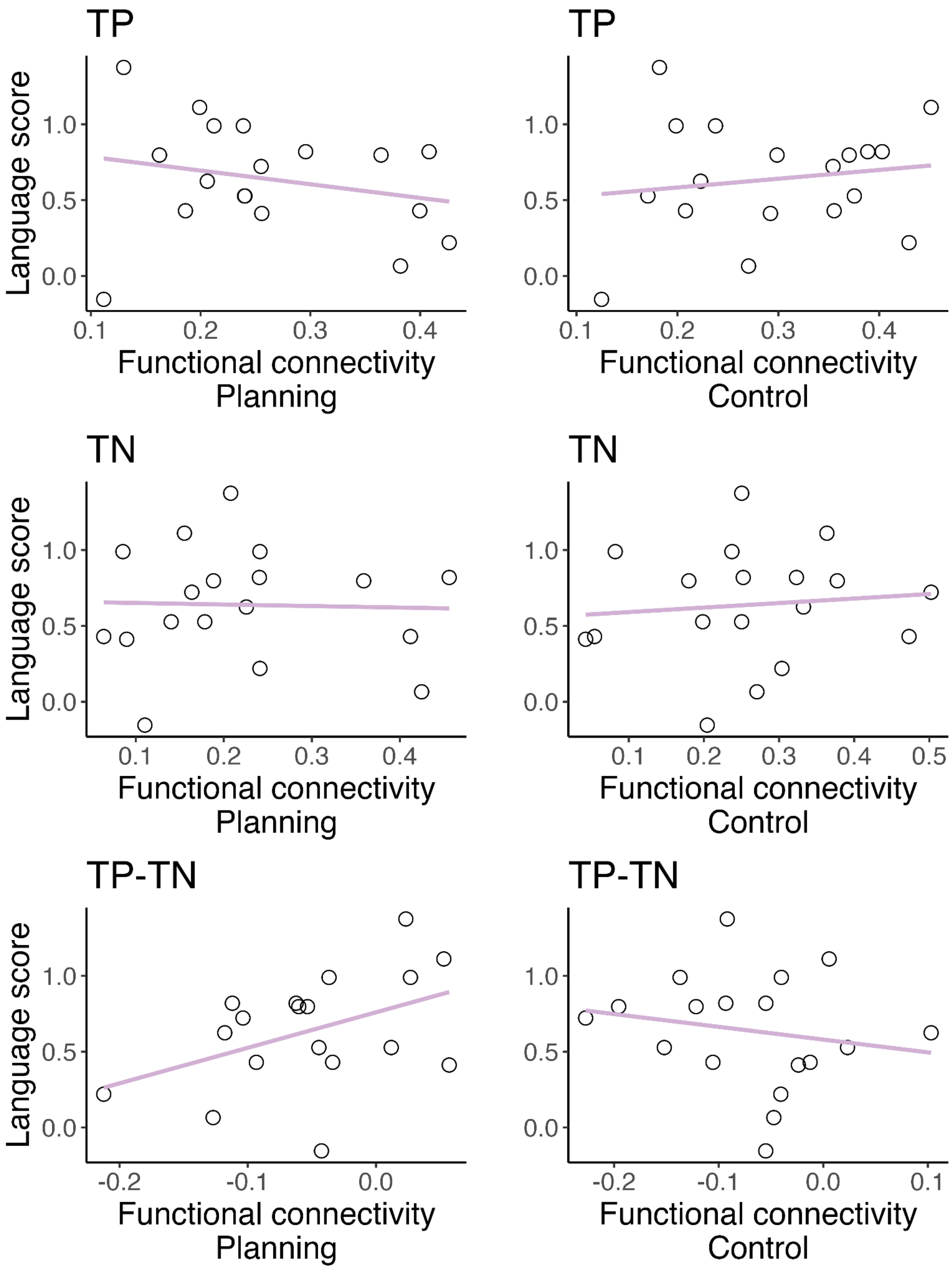

TP
Language score
1.0
0.5
0.0
0.1
0.2
0.3
0.4
Functional connectivity
Planning
TP
1.0
0.5
0.0
0.1
0.2
0.3
0.4
Functional connectivity
Control
TN
Language score
1.0
0.5
0.0
0.1
0.2
0.3
0.4
Functional connectivity
Planning
TN
1.0
0.5
0.0
0.1
0.2
0.3
0.4
0.5
Functional connectivity
Control
TP-TN
Language score
1.0
0.5
0.0
-0.2
-0.1
0.0
Functional connectivity
Planning
TP-TN
1.0
0.5
0.0
-0.2
-0.1
0.0
0.1
Functional connectivity
Control
